# The Impact of Reward on Attention in Schizophrenia

**DOI:** 10.1101/327692

**Authors:** Sonia Bansal, Benjamin M. Robinson, Joy J. Geng, Carly J. Leonard, Britta Hahn, Steven J. Luck, James M. Gold

**Affiliations:** University of Maryland School of Medicine, Maryland Psychiatric Research Center, 55 Wade Avenue, Catonsville, MD 21228; Center for Mind & Brain and Department of Psychology, University of California, 1 Shields Avenue, Davis, CA 95616; Department of Psychology, University of Colorado, 1200 Larimer Street Denver, CO 80217-3364

**Keywords:** selective attention, reward history, spatial probability, schizophrenia

## Abstract

Traditionally, attention was thought to be directed by either top-down goals or bottom-up salience. Recent studies have shown that the reward history of a stimulus feature also acts as a powerful attentional cue. This is particularly relevant in schizophrenia, which is characterized by motivational and attentional deficits. Here, we examine the impact of reward on selective attention.

Forty-eight people with schizophrenia (PSZ) and 34 non-psychiatric control subject (NCS) discriminated the location of a target dot appearing inside a left circle or right circle. The circles were different colors, one of which was associated with reward via pre-training. In the first 2 blocks, targets were equally likely to appear in the left or right circle. In the last 4 blocks, the target was 75% likely on one side, thus allowing us to separately examine how attention was impacted by reward (color) and probability (location).

PSZ had slower overall reaction times (RTs) than NCS. Both groups showed robust effects of spatial probability and reward history, with faster RTs for the rewarded color and for the more probable location. These effects were similar in PSZ and NCS. Negative symptom severity correlated with overall RT slowing, but there were no correlations between symptoms and reward-associated biasing of attention.

PSZ demonstrated RT slowing but normal reward history and spatial probability-driven RT facilitation. These results are conceptually similar to prior findings showing intact implicit reward effects on response bias, and suggest that implicit processing of reward and probability is intact in PSZ.

## 1. INRODUCTION

Since the earliest accounts of Kraepelin (1919) and Bleuler (1950), abnormalities of attention and motivation have been considered to be central features of schizophrenia. Motivational impairments are implicated in the disability associated with the disorder because difficulties initiating and sustaining goal-directed behavior can undermine educational, vocational, and recreational activities. The psychological and neural processes implicated in motivational impairment remain to be determined. Anhedonia is one candidate mechanism given that people with schizophrenia typically report reductions in pleasure on measures such as the Chapman Social and Physical Anhedonia scales (e.g. Horan, Kring, & Blanchard, 2006). However, despite self-reports of low positive affect and pleasurable experience, people with schizophrenia (PSZ) typically show normative affective ratings when actually experiencing positively valenced stimuli under controlled conditions (e.g. Kring & Moran, 2008; Cohen & Minor, 2010). However, for reasons that remain to be fully understood, it seems that the apparently normal hedonic responses at the subjective level fail to have the expected impact on behavior in PSZ. That is, despite evidence of intact in-the-moment reward experience, PSZ typically show reductions in effortful reward-seeking behavior (e.g. Gard et al. 2014).

A possible mechanism by which past reward history may impact future behavior is by influencing selective attention. One central function of selective attention is to reduce the information overload from a rich sensory environment, by prioritizing relevant sensory inputs for further processing.

Traditionally, it has been thought that this prioritization and selection occurs due to either top-down goals or bottom-up sensory salience (e.g. Corbetta & Shulman, 2002). In studies of schizophrenia, selective attention driven by bottom-up information is often unimpaired, whereas deficits in goal-driven control of attention are more frequently reported (Gold et. al, 2007, Luck & Gold, 2008). Recent research has reported a hybrid form of attentional control that may be of special clinical relevance: stimulus selection history (Awh et al. 2012). Stimulus features that become associated with the receipt of rewards in one context may automatically receive preferential processing in a different context where no rewards are available (Anderson et al. 2011; Hickey & van Zoest 2013). This serves an important adaptive function by facilitating approach behavior towards stimuli that have a prior history of being rewarding. People also demonstrate attentional biases based on probabilistic information, even if they are unaware of the probabilities. For example, if a target appears more often at one of multiple potential locations, an attentional bias is likely to develop towards the more frequent location (Geng & Behrmann 2005; Walthew & Gilchrist 2006; Jiang et al. 2013a). Like reward history effects, spatial probabilities are learned quickly, are persistent (Geng et al., 2013; Jiang et al. 2013a; Jiang et al. 2013b) and are typically implicit (Geng & Behrmann, 2005). These effects are top-down insofar as they reflect learning (Gaspelin & Luck, submitted), but they are fast, involuntary, and unconscious, just like bottom-up effects (Theeuwes, 2018).

In order to investigate how attentional control may be differentially impacted in PSZ, we examined the consequences of attentional capture by known reward associations when an implicit spatial probability was introduced. In doing so, we explored the possibility that that PSZ would show reductions in reward history effects as a function of motivational deficits. Although previous studies have exclusively examined aspects of reward processing and spatial allocation of attention in PSZ, it has yet to be understood how multiple sources of selection history interact when presented simultaneously. For example, it may be easier to attend to something associated with pleasure if it is also situated in a predictable location. Outside the laboratory, a reduction in the bias to attend to features and cues associated with past rewards might undermine the initiation of volitional, reward-seeking behavior. This would also be consistent with the idea that PSZ have intact in-the-moment experiences of rewards but that these experiences do not impact the later initiation of reward-seeking behavior (Gold et al. 2009). Alternatively, based on prior studies showing intact implicit reward processing and selective attention in schizophrenia (Erickson et al. 2014; Heerey et al. 2008; Gold et al. 2009; Elshaikh et al. 2015; Barch et al., 2017), another possibility is that PSZ could manifest intact sensitivity to rewards in this task. Under this hypothesis, we expected that PSZ would show intact spatial probability effects because they often show normal levels of benefit from precues that are reliably predictive of target location. In terms of the interplay between the forms of attentional bias, we speculated that if PSZ were impaired at using reward to control attention but unimpaired at using probability, then probability should win when placed in competition with low reward. The paradigm we adapted from Stankevich and Geng (2014), allowed us to examine the relationship between these two factors, and to ascertain our speculations about reward modulation in PSZ.

## 2. METHODS

### 2.1. Participants

Demographic information is provided in Table 1. Forty-eight PSZ were recruited through the Outpatient Research Program at the Maryland Psychiatric Research Center and evaluated during a period of clinical stability (defined as no change in medication type or dosage for four weeks or longer). Consensus diagnosis was established via detailed psychiatric history and interviews, confirmed using the Structured Clinical Interview for DSM-IV (SCID) (First 1995). In PSZ, symptom assessments included the Brief Psychiatric Rating Scale (BPRS, Overall & Gorham 1962) and Scale for the Assessment of Negative Symptoms (SANS, Andreasen 1983).

**Table 1:** Participant Characteristics

|  | HCS<br>(N=34) | PSZ<br>(N=48) | Statistic | p value |
| --- | --- | --- | --- | --- |
| **Age** | 38.53 (11.44) | 38.38 (9.58) | $t = 0.07$ | 0.95 |
| **Gender (M \| F)** | 19 \| 15 | 30 \| 18 | $\varphi = 0.36$ | 0.55 |
| **Race (African American \| Caucasian \| Other)** | 14 \| 18 \| 2 | 16 \| 28 \| 4 | $\varphi = 0.60$ | 0.74 |
| **Participant Education** | 14.88 (2.06) | 13.21 (2.44) | $t = 3.27$ | 0.002 |
| **Maternal Education** | 14.21 (2.56) | 14.21 (2.84) | $t = -0.004$ | 0.99 |
| **Paternal Education** | 15.03 (3.92) | 14.82 (3.59) | $t = 0.87$ | 0.39 |
| **Neurocognitive Test Results** |  |  |  |  |
| WASI-II IQ | 109.44 (11.23) | 92.54 (28.7) | $t = 3.25$ | <0.001 |
| WRAT 4 | 108.65 (14.1) | 93.54 (29.74) | $t = 2.74$ | 0.01 |
| WTAR | 110.68 (13.45) | 94.17 (30.73) | $t = 2.93$ | <0.001 |
| MD Processing Speed | 52.09 (11.66) | 42.76 (11.41) | $t = 3.58$ | <0.001 |
| MD Attention Vigilance | 50.68 (10.54) | 42.26 (12.18) | $t = 3.23$ | <0.001 |
| MD Working Memory | 50.71 (10.61) | 41.85 (11.18) | $t = 3.58$ | <0.001 |
| MD Verbal Learning | 49.35 (8.41) | 38.5 (8.28) | $t = 5.76$ | <0.001 |
| MD Visual Learning | 44.97 (11.02) | 38.85 (10.39) | $t = 2.54$ | 0.01 |
| MD Reasoning | 50.91 (9.93) | 44.02 (10.02) | $t = 3.05$ | <0.001 |
| MD Social Cognition | 55.26 (6.59) | 42.74 (11.16) | $t = 5.83$ | <0.001 |
| MCT Overall | 50.5 (9.92) | 36.11 (12.18) | $t = 5.64$ | <0.001 |
| **Antipsychotic Medication** |  |  |  |  |
| Total CPZ |  | 510.76 (290.21) |  |  |
| Total Haloperidol |  | 10.73 (6.53) |  |  |
| **Clinical Ratings** |  |  |  |  |
| BPRS Positive |  | 2.05 (1.16) |  |  |
| BPRS Negative |  | 1.87 (0.61) |  |  |
| BPRS Disorganization |  | 1.25 (0.32) |  |  |
| BPRS Total |  | 33.39 (11.26) |  |  |
| SANS AA |  | 21.03 (8.55) |  |  |
| SANS EE |  | 13.7 (8.62) |  |  |
| SANS Total |  | 27.15 (10.94) |  |  |

WASI= Wechsler Abbreviated Scale of Intelligence; WRAT = Wide Range Achievement Test; WTAR= Wechsler Test of Adult Reading; MD= MCCB (MATRICS Consensus Cognitive Battery) Cognitive Domain; MCT= MCCB Composite Total; CPZ= Chlorpromazine equivalent; BPRS= Brief Psychiatric Rating Scale; SANS= Scale for the Assessment of Negative Symptoms; AA= Apathy- Avolition; EE= Emotional Expressivity

We also recruited 34 non-psychiatric control subjects (NCS) with no history of psychiatric or substance abuse disorder and no first-degree relative with mental illness, and were matched to the PSZ group for age, gender and parental education. They were recruited by advertisements posted on the Internet and in local libraries and businesses.

Participants were naïve to the purpose of the study and were compensated for participation, all providing informed consent for a protocol approved by the University of Maryland Institutional Review Board. They received an additional monetary payout based on performance. (Mean performance-based earnings: PSZ = $7.44, NCS = $7.60).

### 2.2. Stimuli and Procedure. (See supplemental materials for more details)

The main target discrimination task was preceded by a pre-training task (24 trials) in which participants were acquainted with the task and was designed to teach the participants the reward value of the two colors (Figure 1A).

**Figure 1.**
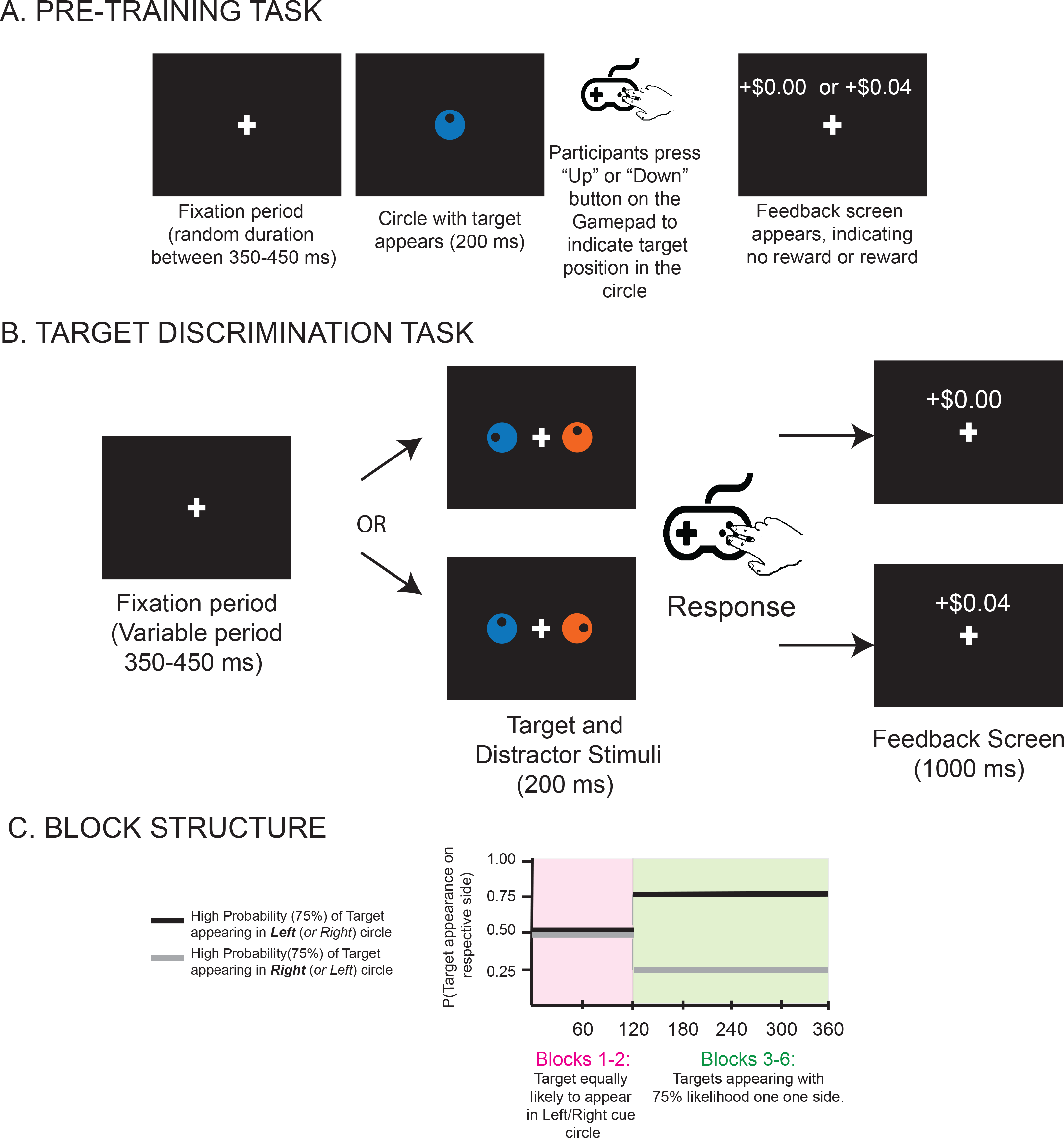
A. Pre-training Task Procedure. A single centrally placed fixation cue (“+”) was presented, after which the cue circle (either blue or orange, rewarded or unrewarded) with the target appeared. **B. Illustration of the trial procedure for the target discrimination task.** Circles are shown in orange and blue to illustrate unrewarded and rewarded colors, respectively. Trials began with a fixation cross, followed by the cue and target screen. Participants were to report whether the target dot was in the top or bottom half of the circle by pressing one of two buttons as quickly as possible. **C. Parametric manipulation of spatial probability.** Participants were assigned to one of two target side high-probability groups, right or left. The staircase shown indicates the course of the task through blocks where the target becomes more likely to fall to one side.

#### 2.2.1. Target Discrimination Task

(Figure 1B). In this task, participants discriminated the location of a target dot appearing inside a left circle or right circle. The circles were different colors, one of which was associated with reward via the pre-training phase. In the first 2 blocks, targets were equally likely to appear in either the left or right circle (equiprobable blocks). In the last 4 blocks, the target was 75% likely on one side (Probabilistic blocks), thus allowing us to separately examine how attention was impacted by reward (color) and probability (location).

### 2.3. Analysis

We included correct trials in which the reaction time (RT) was between 300 and 2000 ms. In the Equiprobable blocks, median RTs were analyzed in a Group by Reward ANOVA, with Group (PSZ vs. NCS) being a between-subject factor, and Reward (unrewarded vs. rewarded) being a within-subject factor. An additional factor of Spatial probability was added for the analysis of the Probabilistic blocks.

## 3. RESULTS

Table 2 contains the statistical test results, including *p* values and effect sizes. Consequently, these statistics will not be provided in the text.

**Table 2:** Statistics

| **A. Equal probability blocks : Collapsed over all trials Group by Reward ANOVA [Fig 2A(i) displays RTs]** |  |  |  |
| --- | --- | --- | --- |
|  | **F** | **p** | **$\eta^2_p$** |
| (i) Reward | 46.84 | < .001** | 0.37 |
| (ii) Group | 7.2 | 0.009* | 0.16 |
| (iii) Reward X Group | 3.55 | 0.076 | 0.04 |
| **B . Equal probability blocks Learning: Learning Rate (from Power Function) ANOVA [Fig 3A]** |  |  |  |
|  | **F** | **p** | **$\eta^2_p$** |
| (i) Reward | 23.98 | < .001** | 0.23 |
| (ii) Group | 1.37 | 0.25 | 0.01 |
| (iii) Reward X Group | 0.36 | 0.41 | 0.004 |
| **C . Probabilistic Blocks Learning (Block 3 & 4): Learning Rate (from Power Function) ANOVA [Fig 3B]** |  |  |  |
|  | **F** | **p** | **$\eta^2_p$** |
| (i) Reward | 378.54 | < .001** | 0.83 |
| (ii) Group | 2.64 | 0.12 | 0.03 |
| (iii) Reward X Group | 0.96 | 0.33 | 0.01 |
| **D. Probabilistic Block [Fig 2(ii) displays RTs]** |  |  |  |
|  | **F** | **p** | **$\eta^2_p$** |
| (i) Group | 13.12 | < .001** | 0.14 |
| (ii) Reward | 58.64 | < .001** | 0.42 |
| (iii) Probability Location | 219.72 | < .001** | 0.49 |
| (iv) Probability Location X Reward | 0.04 | 0.82 | 0.01 |
| (v) Reward X Group | 0.96 | 0.33 | 0.03 |
| (vi) Probability Location X Group | 3.46 | 0.07 | 0.05 |
| (vii) Reward X Probability Location X Group | 0.51 | 0.48 | 0.01 |
| **(viii) Trial Type Comparison [Green arrows, Figure 2(ii)]** |  |  |  |
| Trial Type | 0.11 | 0.74 | 0.001 |
| Group | 18.22 | <0.001** | 0.19 |
| Trial type X Group | 0.02 | 0.9 | <0.001 |

\*\* Significant at <0.001 \*Significant at <0.01

**Table 3.** Correlations [r (p-value)]

|  | SANS Apathy-Avolition | SANS Emotional<br>Expressivity | SANS Total | BPRS Total |
| --- | --- | --- | --- | --- |
| **Overall (across all trial types) Reaction Time** | **0.41**<br>**(0.004)\*\*** | 0.16<br>(0.28) | **0.36**<br>**(0.01)\*** | 0.13<br>(0.27) |
| **Overall Unrewarded- Rewarded RT Difference Score** | 0.004<br>(0.97) | 0.03<br>(0.80) | 0.04<br>(0.78) | -0.16<br>(0.29) |
| **Overall Low- High Probability RT Difference Score** | 0.12<br>(0.43) | 0.28<br>(0.06) | 0.23<br>(0.12) | 0.22<br>(0.15) |

\*\* Significant at <0.01 \*Significant at <0.05

### 3.1. Reward Effects in Equiprobable Blocks

Figure 2A displays the RTs for the first two blocks, (Equiprobable blocks). In these blocks, both NCS and PSZ had slower RTs when the target appeared in the unrewarded colored circle than when it appeared in the rewarded colored circle (significant main effect of reward) with PSZ being slower overall than NCS (significant main effect of group). NCS exhibited a slightly larger effect of reward than PSZ, but the interaction between Group and Reward did not reach significance (Table 2A. (i-iii)). Note that the effect size was large for the main effect of Reward (η^2^_p_ = 0.37) but small for the Group X Reward interaction (η^2^_p_ = 0.04). We also performed comparisons of the rewarded and unrewarded colors separately for each group, and we found a large and significant Reward effect in both PSZ (t=4.01, p<0.001, Cohen’s d = 0.58) and NCS (t=5.54, p<0.001, Cohen’s d = 0.95). Thus, although the effect of reward was numerically larger in NCS than in PSZ, reward had a large and significant effect in both groups and the differences in the reward effect between groups were not significant.

**Figure 2.**
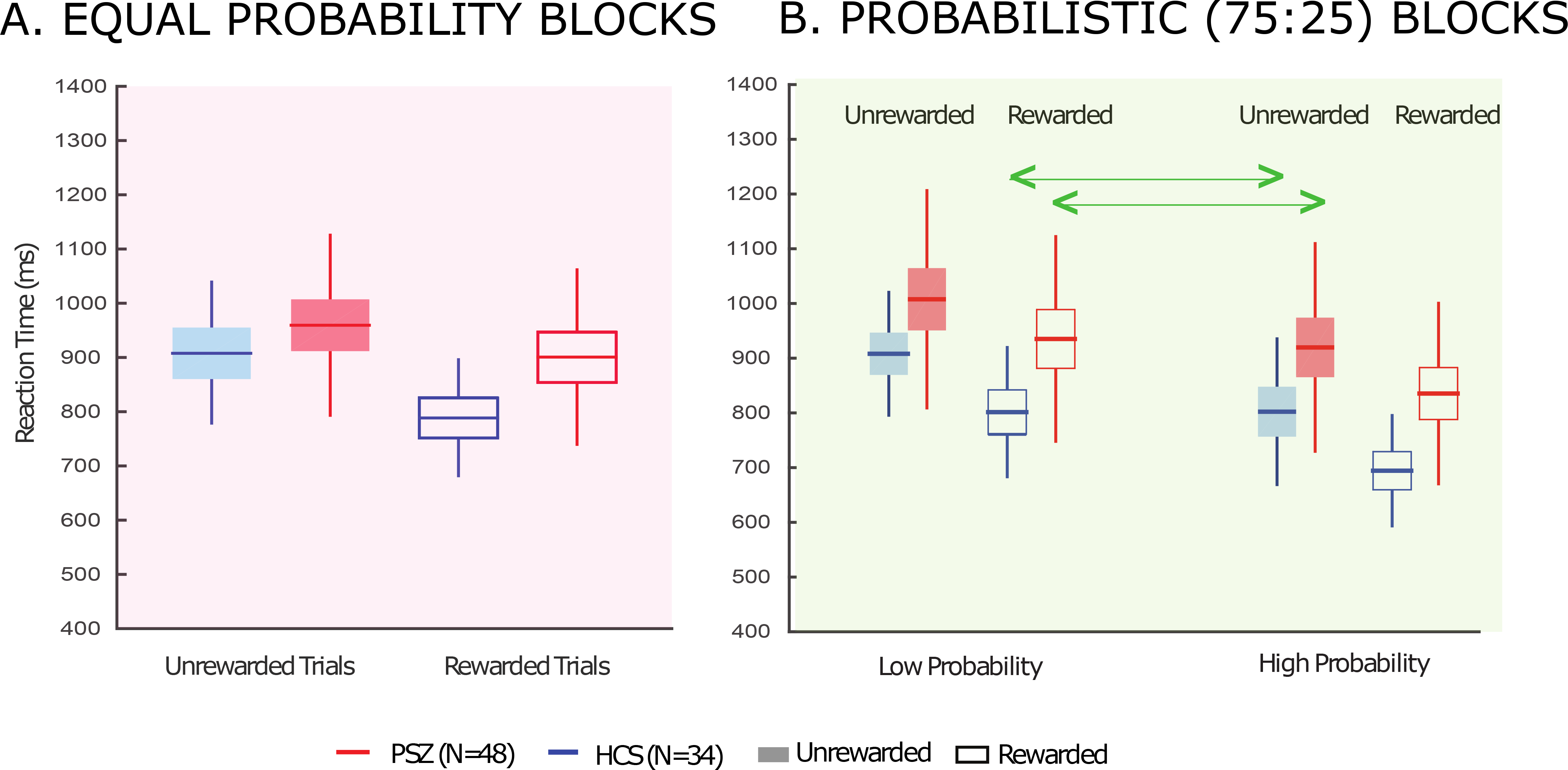
Median RT results for Target Discrimination Task. **A.** Average of Median RTs for unrewarded and rewarded trials at for equal probability blocks. Reaction times in both groups were shorter to rewarded than to non-rewarded targets. **B.** Average of Median RTs for unrewarded and rewarded trials for low probability and high probability trials respectively. Reaction times in both groups were shorter to rewarded than to non-rewarded targets for both probabilities, and an increase in the strength of the probability resulted in shorter RTs. The arrows indicate the trial types for which the two factors, spatial probability and reward were in conflict.

Further, both groups learned the color-reward association at a similar rate. Figure 3 shows how RTs changed over the first two blocks in the main target discrimination task (following the pre-training phase) for the rewarded and unrewarded colors, binned into 6 equal bins of 10 trials each for display purposes. We fit a power function to RTs over all trials in each subject, which provides the standard metric of learning rates in RT experiments (Logan, 1988; 1992). A Group by Reward ANOVA (Table 2B) on individual learning rates yielded no significant Group by Reward interaction effect, nor a main effect of Group, with near-zero effect sizes. However, the main effect of Reward was significant, with a large effect size. Note that by the last time bin, the difference in RT between the rewarded and unrewarded colors was similar in PSZ (129.52±26.52 ms) and NCS (147.87± 31.10 ms).

**Figure 3.**
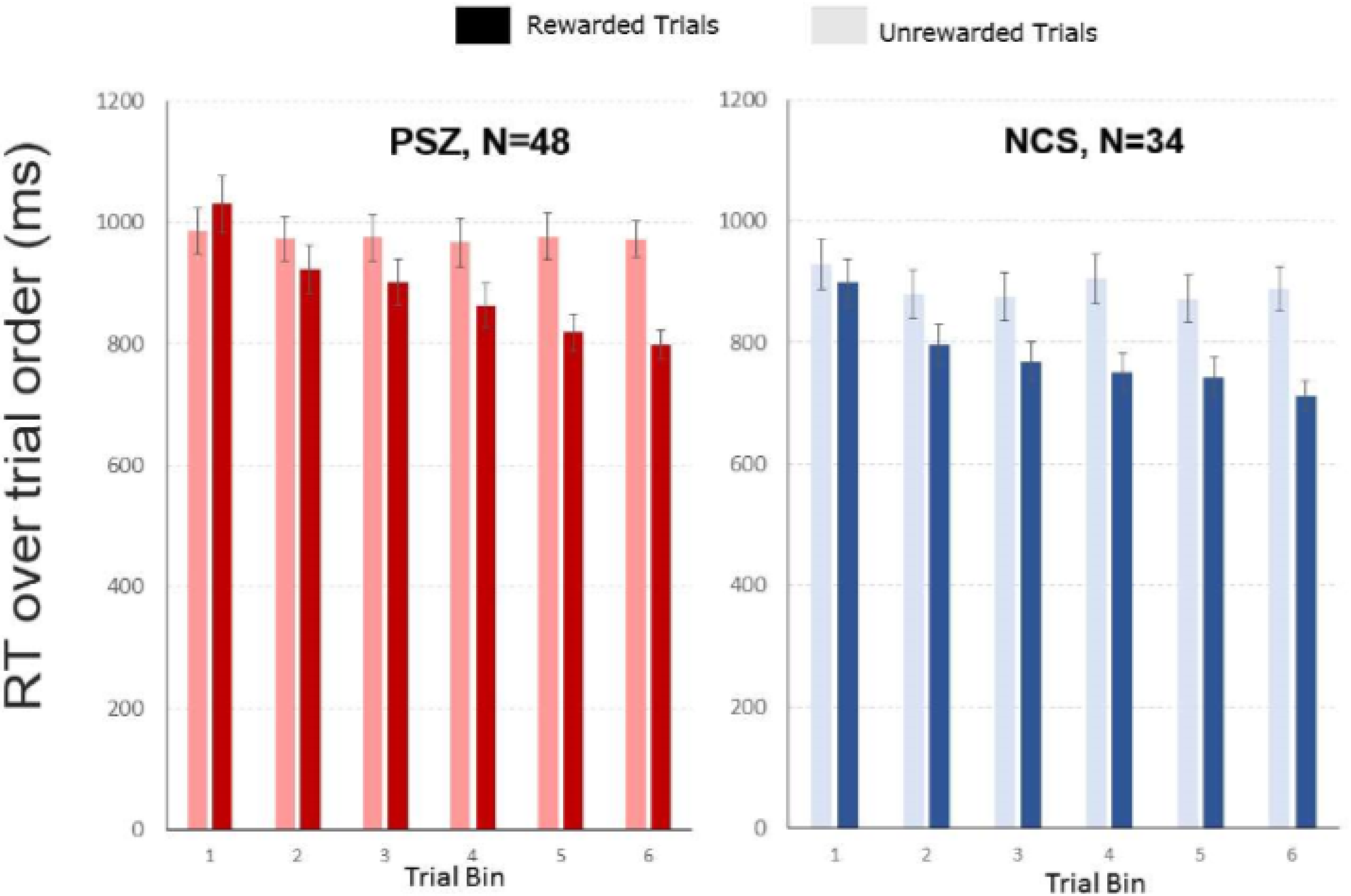
RT speeding over rewarded trials in equal blocks. The y axis displays the mean RTs per trial bin over the first two blocks (equal probability blocks). For the x axis, the 60 trials of each participant per condition (Rewarded and Unrewarded) were binned into 6 bins of 10 trials and averaged across participants. The error bars display the standard error of the means in the respective trial bin. Participants showed a significant decrease of RTs for rewarded trials (Darker Shaded bars) over the course of trials, while there was no such decline for unrewarded trials (lighter shaded bars). Red bars represent data from PSZ, while Blue bars represent the same for NCS. This was corroborated by no significant Group by Reward interaction for learning rates.

### 3.2. Reward and Probability Effects in Probabilistic Blocks

Similar to the learning rate analysis conducted for reward effects, we fitted a power function to the RTs over high-probability and low-probability trials (independent of reward) in the first two probabilistic blocks to examine probability-location association learning. A Group by Probability ANOVA on these learning rates yielded no significant Group by Probability interaction effect, nor a main effect of Group. (Table 2C)

Figure 2B displays the RTs for the Probabilistic blocks as a function of reward and spatial probability, with statistical results indicated in Table 2D. We found that: (i) PSZ had slower reaction times overall; (ii) RTs in both groups were shorter to targets appearing in the rewarded circle than in the unrewarded circle; (iii) RTs in both groups were shorter to targets appearing in the high-probability location than in the low-probability location; and (iv) the effects of reward and probability were largely independent, with a similar effect of reward at the two locations. A similar independence of reward and probability effects was found in a study of healthy college students (Stankevich & Geng 2014).

These impressions were confirmed statistically by significant main effects of Location and Group. The Reward by Group interaction did not approach significance, with a very small effect size. The speeding of RTs at the high probability location relative to the low probability location was slightly greater in NCS than in PSZ, but the Location by Group interaction did not reach significance and the effect size was small. We did not observe any other 2- or 3-way interactions involving reward that approached significance. We also compared rewarded and unrewarded trials (collapsed across probability) for each group separately, and we found a large and significant Reward effect in both PSZ (t=4.90, p<0.001 Cohen’s d = 0.71) and NCS (t=6.86, p<0.001, Cohen’s d = 1.17). We also compared the low and high probability locations (collapsed across reward) for each group, and we found a large and significant Probability effect in both PSZ (t=7.01, p<0.001 Cohen’s d = 1.01) and NCS (t=11.37, p<0.001 Cohen’s d = 1.95). Thus, although RTs were slowed overall in PSZ, both PSZ and NCS showed robust effects of both reward history and spatial probability, and group differences in these effects were not significant and had small effect sizes. The combination of large effect sizes for the Reward and Location main effects and the small effect sizes for the Reward by Group and Location by Group interactions provides compelling evidence that any differences between groups in reward and probability effects are small.

The attentional benefit conferred to the target at the higher probability location did not change as a function of the target’s reward incentive, and neither was the benefit of the rewarded color altered by the target’s location, as indicated by a near-zero effect size for the Location by Reward interaction. This indicates that, in accordance with the findings of Stankevich & Geng (2014), these two sources of information that guide selective attention act independently and have an additive effect. We chose spatial probabilities of 75% and 25% for the Probabilistic block because Stankevich & Geng (2014) found that the effect of probability was approximately equal to the effect of reward in this paradigm for these probabilities, leading to approximately equal RTs when these two factors conflicted with each other (i.e., on rewarded low-probability trials and unrewarded high-probability trials, indicated by green arrows in Figure 2). If PSZ were impaired at using reward to control attention but unimpaired at using probability, then probability should win when placed in competition with low reward, leading to faster RTs for unrewarded high-probability trials than for rewarded low-probability trials. However, both PSZ and NCS in the present study also exhibited nearly equal RTs for these two trial types. This observation was supported by a separate 2-way ANOVA (Table 2D (i)) comparing these two trial types in PSZ and NCS (in the Probabilistic blocks), which yielded no significant main effect of trial type and no significant interaction between group and trial type. These results are consistent with the hypothesis that PSZ are not impaired at using reward information to control attention.

Taken together, these results show that PSZ and NCS were similarly impacted by reward during the early blocks, and even if the effects of reward and spatial probability in the two groups over all the experiment blocks are not the same, the effect size statistics indicate that group differences are subtle.

### 3.3. Correlations

To determine how performance in the present task was related to symptoms, we examined correlations between median RTs in both tasks and BPRS and SANS scores. We observed significant correlations between overall RT and SANS scores (Avolition-Apathy: r=0.41, p=0.004; SANS Total: r=0.36, p=0.01. Thus, overall slowing of responses was associated with increased levels of negative symptoms.

There were no correlations between SANS scores and the effects of reward (unrewarded RT minus rewarded RT), nor between SANS scores and effects of probability (low minus high probability). Neither reward effects, nor probability effects were associated with positive symptoms. Thus, overall slowing of RT was correlated with increased negative symptoms, but impairments in reward effects and probability effects were not.

## 4. DISCUSSION

To our knowledge, this experiment is the first to examine the role of learning on selective attention in people with schizophrenia. We explored two different varieties of learning: the ability to associate a reward value with a specific color and the ability to associate a target stimulus with a specific location. Somewhat surprisingly, we found that both aspects of attentional control were relatively intact in people with schizophrenia. For reward value, we found that both PSZ and NCS exhibited significant effects of reward in both the Equiprobable and Probabilistic blocks (Cohen’s d > 0.58), with the effect sizes for the Group by Reward interaction being small (η^2^_p_ < 0.04), indicating that this interaction accounted for less than 4% of the explainable variance in RTs. In contrast, the effect sizes were larger (η^2^_p_ > 0.08) for the main effects of group, reward, and probability. Moreover, learning rates over the first two blocks of trials were similar for PSZ and NCS, with a near-zero effect size for the Group by Reward interaction. We found a similar pattern for the probability effect, with large and significant effects of probability in both PSZ and NCS (d > 1.01), a large effect size for the main effect of Probability (η^2^_p_ = 0.49), and a small effect size for the Group by Probability interaction (η^2^_p_ = 0.05). Although we are not claiming that the effect of reward and probability on attention are the same in PSZ and NCS, the present results provide substantial evidence that both PSZ and NCS are highly sensitive to both reward and probability, and any differences between groups are small (and presumably clinically negligible).

It is important to consider the possibility that the PSZ and NCS samples in the present study were non-representative, with either unusually high-performing PSZ or unusually low-performing NCS. This is unlikely given that our PSZ sample was strongly impaired relative to NCS on neuropsychological measures such as IQ (see Table 1). In addition, the PSZ showed the typical elevation of overall RTs relative to NCS, and the degree of RT slowing was significantly correlated with motivation-related negative symptoms, which in contrast, were not associated with reduced reward or probability effects. Motivational deficits as assessed by clinical assessments were not specifically associated with the modulation of attentional capture by reward and probability.

The present study provided evidence that patients could utilize both reward associations and spatial probability to enhance the speed of their reaction times, as evidenced by similar learning rates. Thus, even though impairments of attention are often claimed to be central features of the disorder, many aspects of selective attention appeared to be intact in the present experiment. This is consistent with other recent studies using methods that could isolate selective attention from other cognitive processes (e.g. Erickson et al. 2014; Elshaikh et al. 2015). Such areas of intact function may be of particular interest because they argue against the claim that PSZ have a “generalized deficit”, impacting nearly all aspects of cognitive function. That generalized deficit does not extend to important aspects of selective attention as seen here.

The finding that reward-based control of attention is intact in PSZ clearly contrasts with other aspects of reward processing that are impaired in schizophrenia, such as effort-based decision-making and rapid reinforcement learning (e.g. Barch et al., 2017; Strauss et al. 2014; Fervaha et al. 2013; Barch et al. 2014; Gold et al. 2012; Heerey et al. 2007; Barch & Dowd 2010). Impairments are most likely to be observed when representations of expected value are needed to guide decisions or when rapid behavioral adjustments are needed in response to feedback (Gold et al. 2008; Gold et al. 2012). The combination of the present study and previous research (see reviews, Hélie et al. 2017; Strauss et al. 2014) provides evidence that the binary evaluation of outcomes (reward versus non-reward) may be intact in schizophrenia, but other aspects of reward processing—such as the effect of outcome magnitude, weighing costs versus benefits of effortful responding, and encoding of expected reward value (Strauss et al. 2014; Gold et al. 2013; Waltz et al. 2009)—may be impaired. It appears likely that motivational deficits are more closely related to these complex aspects of reward processing and decision making than to implicit biasing of attention.

Areas of intact implicit reward processing are important because they indicate that avolition in schizophrenia is not a result of insensitivity to rewards. Indeed, intact implicit reward processing may be an important lever that could be used in developing psychosocial interventions to enhance motivational functioning. From a basic cognitive neuroscience perspective, the results support previous findings (Stankevich & Geng 2014) in showing that reward associations and spatial probabilities are separate but powerful cues that guide attentional selection. Interestingly, these factors had independent effects across both college student populations in the previous study and a broad range of people with and without schizophrenia in the present study. In the decision-making literature, these factors are typically combined into a single metric combination of probability and reward value (i.e., expected value); for the purpose of attentional selection, however, these two factors act separately and produce additive behavioral effects.

A limitation of the present study is that—even though we provided actual monetary rewards and successfully replicated basic cognitive neuroscience findings—the external validity of the present task is unknown, and results may not translate to real-world reward history-based selective attention and perceptual processing. Also, only stable outpatients receiving antipsychotic therapy were included (although we found no significant correlation between chlorpromazine equivalents and any of our RT measures, r <0.25, p>0.17). Similar work with more severe inpatients or unmedicated or medication-naïve populations may yield different results. However, inclusion of stable outpatients enabled us to examine participants with generally lower levels of acute symptoms, allowing the feasibility of measures involving self-report and computerized testing.

In conclusion, we replicated previous findings showing that reward history and spatial probability modulate selective attention and have additive, independent effects. We further provided evidence that this modulation is similar in PSZ and NCS, thus adding to the literature suggesting that implicit reward processing in the context of visual selective attention and target discrimination may be surprisingly intact in PSZ.

## Contributors

Sonia Bansal undertook the statistical analysis and interpretation and wrote the first draft of the paper. Benjamin Robinson contributed to recruitment of participants, data collection and reviewed the manuscript. Joy Geng, Steven Luck and James Gold designed the study and wrote the protocol. Carly Leonard and Britta Hahn contributed to study design and critical review of the manuscript. Steven Luck and James Gold developed the study, raised funding, and contributed substantially to analysis and interpretation and the writing of the manuscript. All authors contributed to and have given final approval of the manuscript.

